# Explainable AI-guided identification of a novel protein-RNA interactive frame for selective siRNA accumulation in plants

**DOI:** 10.1101/2024.12.02.626406

**Authors:** Natsumi Enoki, Eriko Kuwada, Shinnosuke Matsuo, Naoko Fujita, Saki Noda, Yoshikatsu Matsubayashi, Seiichi Uchida, Takashi Akagi

## Abstract

Small interference RNA (siRNA) selectively accumulates and acts in RNA interference (RNAi). Although the components involved in siRNA production have long been the focus of studies to elucidate RNAi processes, the mechanism(s) for selectivity of siRNA (or RNAi effectivity) remains unclear. In a novel approach, we developed a progressive deep learning (DL) framework integrating Transformer and convolutional neural networks to predict the sequences of selectively accumulated siRNAs across various land plant species. These approaches achieved high-accuracy prediction of selectively accumulated 21-nt siRNAs and further identified their key signals, which are positionally and linguistically flexible sequences surrounding the target siRNA. We experimentally validated the contribution of these flexible key signal sequences to siRNA accumulation selectivity using virus-induced gene silencing (VIGS) in *Nicotiana benthamiana*, and identified RNA-binding proteins that directly recognize the key signal sequences to act for selective siRNA accumulation. These insights provide a novel framework for investigating RNAi mechanisms in plants.

**One sentence summary:** We discovered a novel mechanism centered on RNA-protein interactions involving the selective accumulation of small-RNAs in plants by applying advanced deep learning frames.

## Introduction

In plants, RNAi is a common and fundamental system for regulating gene expression that is based on double-stranded RNA-dependent post-transcriptional gene silencing^1^. Small interference RNAs (siRNAs; 21-24 nt), which are derived from the cleavage of dsRNA by Dicer-like 1-5 (DCL1-5) in plants, play a main role in this gene silencing process^2^. After siRNAs (or guide siRNAs) are produced, they, especially *trans*-acting 21-nt siRNAs and phased 21-nt siRNAs, are selectively recruited to the target mRNA. They effectively degrade the target mRNA and often stimulate the biogenesis of secondary siRNA as well^3–4^. However, only a limited fraction of siRNAs can be highly accumulated via this selective recruitment or recycling process, which effectively causes RNAi^3^ (see Fig. 1a, Fig. S1 for *ChlH*). Currently, little is known about the molecular mechanism underlying this process. Previous studies have attempted to identify the nucleotide sequence patterns specific to highly effective 21-nt siRNA and have found some biases in the localization of specific residues in 21-nt siRNAs^5–6^. For instance, the highly effective anti-sense strand siRNA tends to be enriched in A or U in the 5-bp of the 5′ end-sequences, whereas the highly effective sense strand tends to start with G or C at the 5′ end^5^. However, a genome-wide survey suggested that there are abundant exceptions in many plant genomes (Fig. S2 for the residue distribution). Currently, neither the systematic nucleotide sequence patterns nor the genetic factors determining the selectivity of highly accumulated siRNAs have been fully defined.

**Figure 1.**
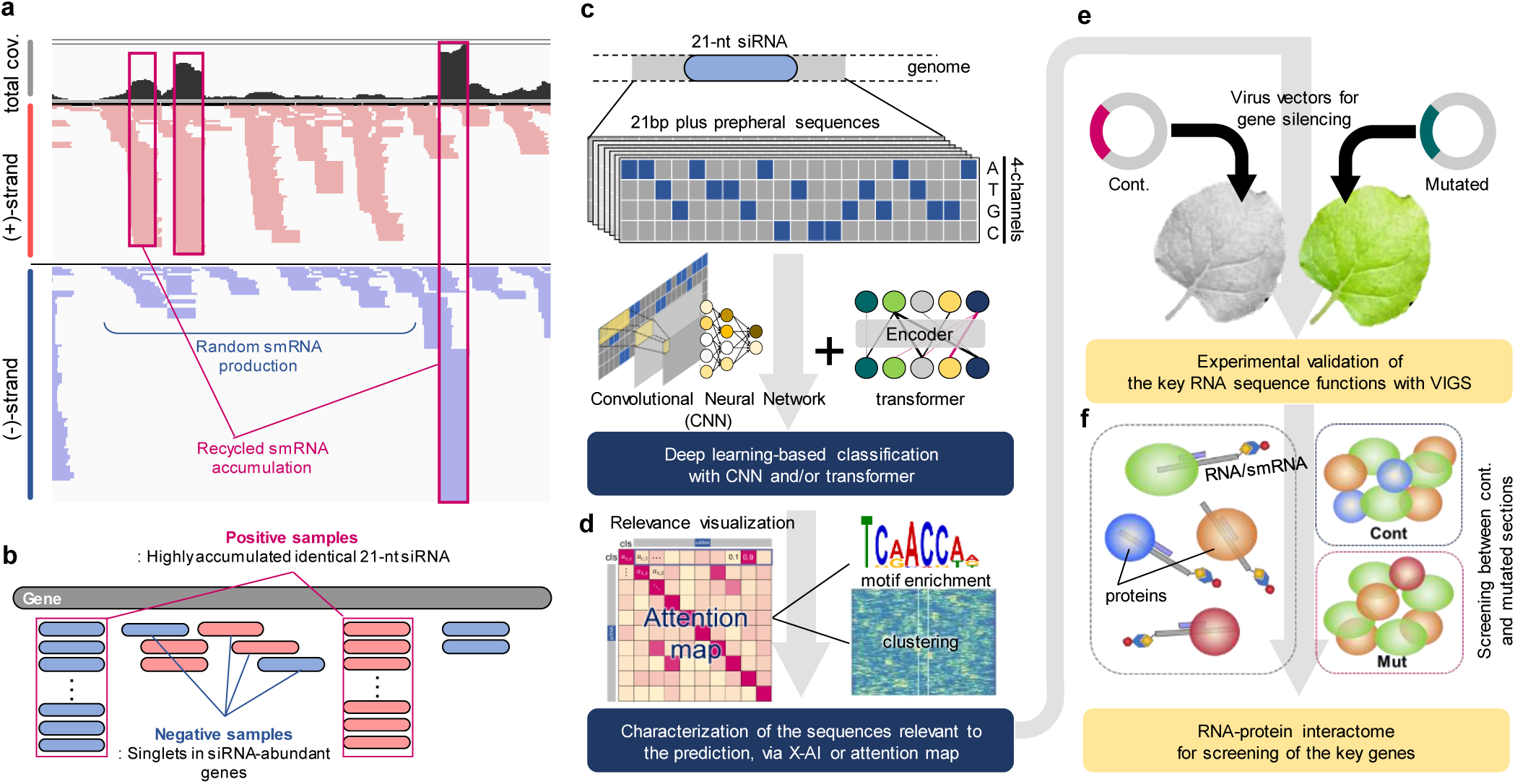
Concept and experimental designs of this study. **a**, A small-RNA-seq result indicating that a limited number of 21-nt siRNAs were selectively recycled and highly accumulated in a strand-specific manner. **b**, Sample categories used for the following deep learning analyses. **c-d**, Experimental flow with explainable deep learning (or X-AI) to predict highly accumulated siRNA (**c**) and characterize their features (**d**). (**e-f**), Experimental flow for validating the AI-predicted features (or signal sequences) using virus-induced gene silencing (**e**), and for RNA-protein interactome analysis to identify genetic factors that interact with the AI- predicted signal sequence required for siRNA accumulation (**f**).

In biological contexts, there are some methodologies to identify nucleotide sequence patterns. Position weight matrix (PWM)-based methods, represented by MEME series^7^ or *k*-mer-based machine learning methods^8–9^ have been effective in capturing specific sequences contributing to an objective, such as expression regulation. Recent progress in deep learning has also allowed for capturing more flexible linguistic contexts. The development of Transformer^10^ resulted in a piece of key architecture in many fields. In biology, for instance, Transformer has been applied to the core driver of the AlphaFold series^11–13^, which predicts protein 3D-structures from amino acid sequences. It has also been applied in an expression prediction model for long promoter nucleotide sequences, and its performance in this context has been outstanding^14^. The convolutional neural network (CNN) has been a classical deep learning framework mainly used for image analyses^15–16^, and it can consider features in flexible positions (or with the “translation-invariant” characteristics of CNN). CNN is also applicable to wide data formats, including nucleotide sequences^17–19^. Although these technical progresses would be potentially able to unveil various unsolved nucleotide sequence contexts, there have been a limited number of applications in plant biology. Here, as shown in Fig. 1, we first aimed to define the nucleotide sequence patterns that determine the selectivity for highly accumulated 21-nt siRNA using integrative explainable AI (X-AI) models of Transformer and CNN. Second, we attempted to identify the genetic factor(s) that recognize this signal sequence and could potentially regulate selective siRNA accumulation by interacting with it.

## Results and Discussion

We collected 18 small-RNA (smRNA)-seq datasets from various species/organs, including a wide variety of land plants (see Table S1). They were mapped to each reference genome to define read numbers per gene and smRNA coverages. We defined highly accumulated 21-nt siRNA (cov. >20) and singlets in the siRNA-enriched genes/regions as positive and negative samples, respectively (Fig. 1a-b, see Methods for the details). The 21-nt sequences (U was converted to T) and both of their flanking genomic sequences (0, 15, 40, 60, and 90 bp) were converted into one-hot arrays with four channels (A, T, G, and C)^20^ (Fig. 1c-d, Fig. 2a). We constructed four deep learning frameworks (fully connection, VGG-like 1d-CNN, Transformer encoder, and Transformer/CNN integrative models, Fig. S3 shows the model architectures) to classify the described positive/negative samples. In the prediction model trained with rice (*O. sativa*) smRNA data, the integrative model with CNN and Transformer (Convolutional vision Transformer; CvT) exhibited the best performance among the models examined, and it achieved >0.86 in Receiver Operating Characteristic Area Under Curve; ROC-AUC value (Fig. 2b). Importantly, training with longer flanking sequences of the targeted 21-bp siRNA fundamentally improved the prediction performance in all the models we used (Fig. 2b). These tendencies were conserved in the other smRNA datasets from various species (Fig. 2c). This indicated that the feature characteristics (or key signals) for selective siRNA accumulation are located not only inside the 21-nt siRNA but also in the peripheral regions, which appears to be common among the land plant species. Across the smRNA datasets, the CvT model achieved the best performance (Fig. 2c, the best ROC-AUC = 0.95 with >90% accuracy in the *Arabidopsis thaliana* model). Transfer learning among the species (or interspecific application of the trained models) indicated that the prediction performance was not dependent on phylogenetic closeness (Fig. 2d), implying that a common mechanism for siRNA selectivity is conserved among a wide variety of land plant species, at least to some extent. Note that the prediction ability tended to be highest in the own (trained) species (Fig. 2d). This would be presumably due to the fact that a single signal sequence can be shared amongst multiple highly accumulated siRNA, as discussed later (discussed in Fig. S6). Furthermore, frequent “lineage-specific” gene/genome duplication events in plants, from an aspect of evolutionary biology^21,22^ might be a reason. Although some of the functional properties are adaptively changed in the duplicated regions^23^, most regions still maintain similar genomic properties, particularly in recently duplicated regions. The prediction performance was slightly dependent on the trained sample numbers, in which the accuracy tended to improve until the number of training samples reached 40,000 (Fig. S4a). Consistent with this result, training with the interspecifically merged samples (*N* > 50,000) did not result in an improvement in prediction ability (Fig. S4b).

**Figure 2.**
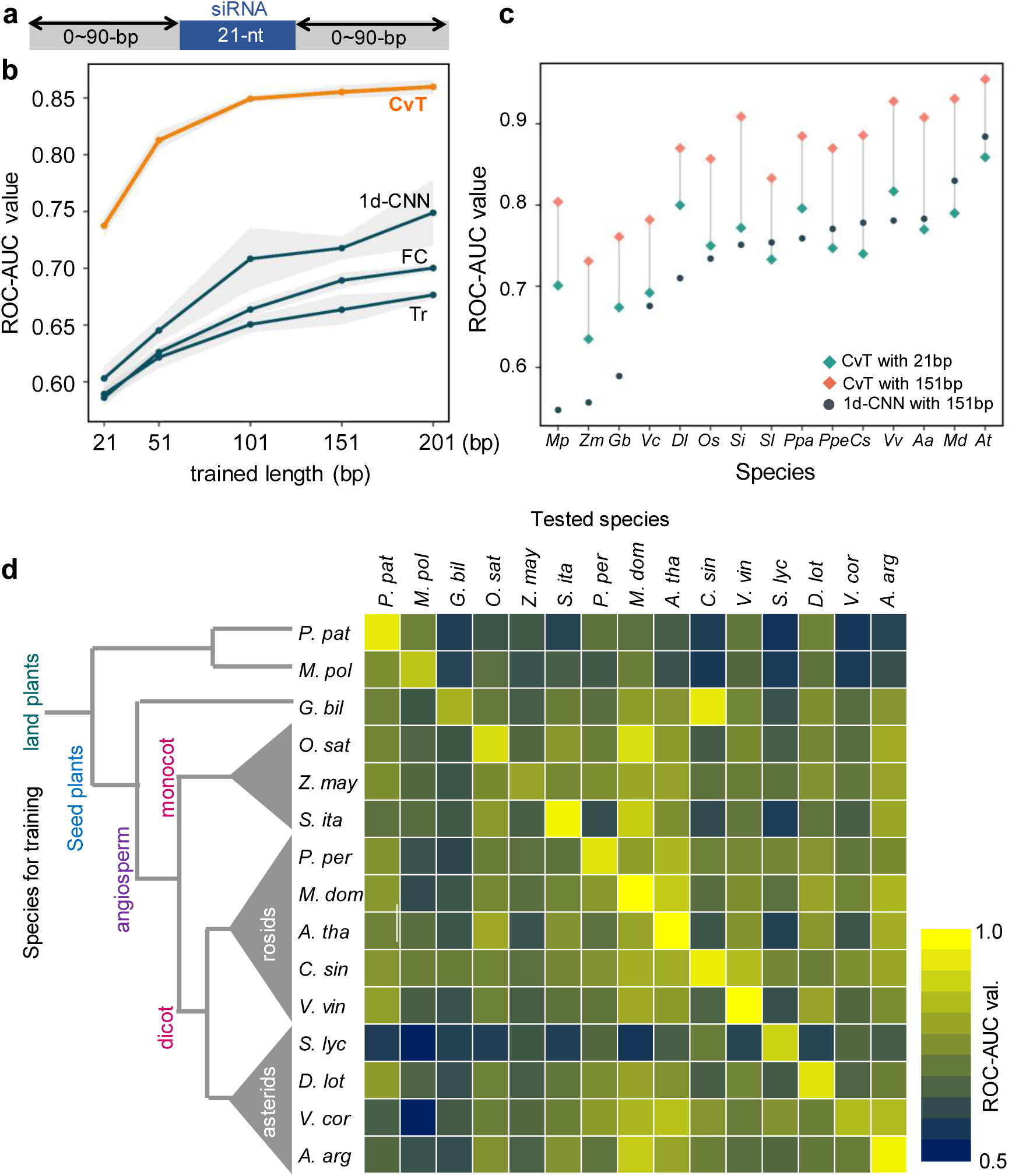
Transformer-CNN integrative deep learning model (CvT) highly predicts selectively accumulated siRNA in a wide variety of land plants. (**a-b**), Performance of four deep learning models trained using different length flanking sequences of the target 21-nt siRNA. (**b**), CvT, CNN-Transformer model, 1d-CNN, one-dimensional CNN model, FC, fully connected model, and Tr, simple Transformer model. The prediction performance was evaluated using ROC-AUC. Training with longer flanking sequences of the targeted 21-bp siRNA fundamentally improved the prediction performance in all the models we used. (**c**), Distribution of the prediction performance (ROC-AUC) after training with smRNA-seq data from various plant species, comparing 21-bp/151-bp for the sequence length and CvT/1d-CNN for the model. For all of the plant species tested, training with longer (151-bp) sequences meant that the CvT model achieved better performance, up to ROC-AUC = 0.955. (**d**) Transfer learning amongst a wide variety of land plant species. The CvT models trained with 151-bp (21-bp siRNA + flanking 65-bp x 2) were applied. The phylogenetic distance was mostly independent of the prediction ability, suggesting that a common mechanism (or sequence patterns) may involve the selective siRNA accumulation amongst the land species.

The positive sample fraction with the highest prediction confidence (conf. > 0.9, *N* = 694) in the *O. sativa* CvT model that was trained with sequences 151-bp (21-bp + 65-bp x 2) in length (Fig. 3a) exhibited no enrichment of specific genomic/functional annotations compared with the whole positive samples (Fig. S5). This supported the conclusion that our deep learning prediction is not affected by specific characteristics of the gene or transposable element but is only dependent on nucleotide sequence patterns. In positive samples with the highest prediction confidence, we visualized weighted attention (or hypothetical features relevant to the prediction in Transformer) (Fig. 3b-c). As expected, the key regions for selective siRNA accumulation are located not only inside the 21-bp siRNA but also in the flanking regions. The attention weight pattern in the 151-bp region indicated no clear tendency toward clustering (Fig. 3c), suggesting that the key region is positionally flexible. By contrast, the weighted attention was common for independently accumulated siRNA close to one another (Fig. S6). This suggests that a single nucleotide sequence can be used as a signal for multiple siRNA accumulations. PWM-based MEME analysis^7^ to search for enriched gap-free motifs was also conducted on the sequences with the highest prediction confidence in the *O. sativa* CvT model (Fig. 3d, *N* = 694). However, even the top-hit enriched motif exhibited low confidence values (e-value = 1.9e^-5^), and it was included in only <1.2% of these sequences. Even the allowance of gaps for the enriched motifs in GLAM2^24^ did not result in any clear and confident patterns (Fig. 3e, e-value = 2.1e^-6^). Similarly, in the other species, the top-hit enriched motifs in the sequences with high prediction confidence in each CvT model explained only <11% of the whole sequences (Fig. 3f). Even the total statistically enriched motifs (e-value < 10) appeared in less than 30% of the whole sequences (Fig. 2f). Additionally, there were no conserved sequence patterns in the enriched motifs among the plant species (Fig. S7). Thus, our CvT model successfully captured the positionally flexible signal sequences contributing to siRNA accumulation, although it might be difficult to express them as simple nucleotide patterns.

**Figure 3.**
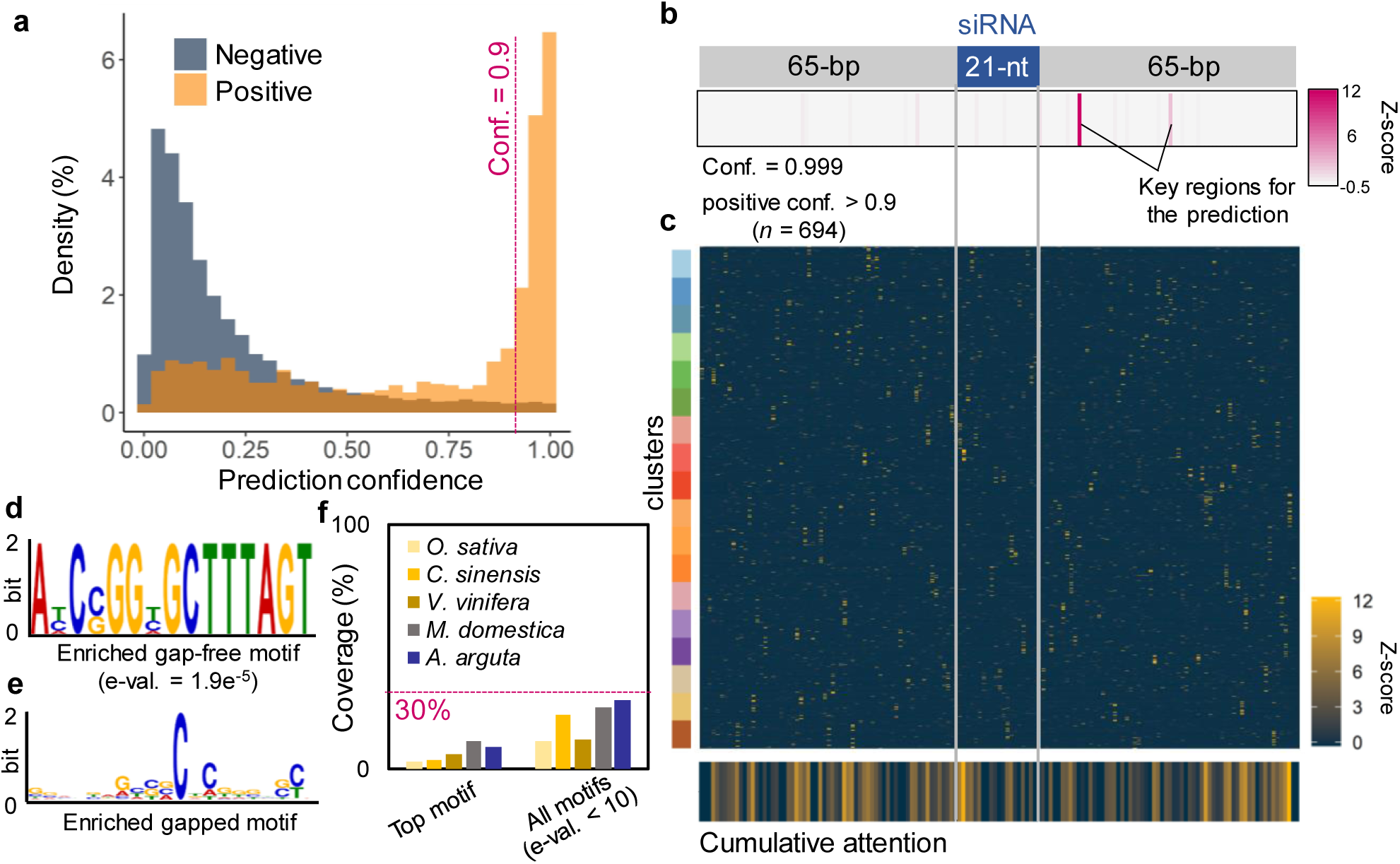
Examination of feature characteristics in the prediction of key signal of selectively accumulated siRNA. (**a**), Distribution of the prediction confidence of actual positive/negative samples in the *O. sativa* CvT model. For the following analyses, we selected the highly confident positive samples (conf. > 0.9 for the positive, *N* = 694) to characterize the features as positive. (**b-c**), Locations of the standardized (Z-scored) weighted attention (or key region for the prediction) in a 151-bp fragment including a positive target siRNA (**b**), and in the whole samples with high prediction confidence (*N* = 694) (**c**). In panel (**c**), the attention distribution was hierarchically clustered, although no tendency was observed. (**d-f**), Motif enrichment analysis in the weighted attention-surrounded regions (21- bp). The top-hit enriched gap-free motif analyzed by MEME (**d**) and gapped motif analyzed by GLAM2 (**e**), in the *O. sativa* CvT model. Explainability with the top-hit enriched motif and with all statistically significant motifs (e-value < 10) in various plant species (**f**).

To experimentally validate the function of the key signal sequences our DL models predicted, we applied VIGS in *Nicotiana benthamiana* using the apple latent spherical virus (ALSV)^25^ (Fig. 1e). *ChlH* (a gene involved in chlorophyll biosynthesis) fragments, which were predicted to cause the high accumulation of siRNA in our *Citrus sinensis* models (prediction conf. > 0.95) were inserted into the ALSV vector, and leaf decoloration was used as the index for determining the level of siRNA accumulation^26^ (Fig. 4a-e). We preliminarily detected some highly accumulated (or positive) 21-nt *ChlH* siRNA in *N. benthamiana*, but the insertion of only these positive 21-nt *ChlH* fragments resulted in no decoloration in the ALSV-infected plants (Fig. 4c). However, when both flanking sequences, including the weighted attention (or key) region (Cont., see Fig. 4b), were inserted, clear decoloration was observed (Fig. 4d). By contrast, when the key regions were point-mutated (Mut., see Fig. 4b), the decoloration effect was mostly lost (Fig. 4e). A small-RNA-seq analysis of these Cont. or Mut.-insert infected *N. benthamiana* leaves showed that when point-mutations in the key regions were introduced (Fig. 4b), 21-nt siRNA peaks were lost not only in the targeted central siRNA but also in the surrounding siRNAs (Fig. 4f-g). This is likely because signal sequences are shared for multiple highly accumulated siRNAs whose sequences are close to one another, as shown in Fig. S6. These findings were not limited to the *ChlH* gene; infection with *PDS*, *AGO1*, and *PHAN* fragments with high prediction confidence for the positive in the CvT/1d-CNN models for various plants resulted in effective gene silencing (Fig. S8). Next, based on a positive 153-bp *ChlH* insertion with high prediction confidence (conf. = 0.999), we defined a minimal insert (51-bp) region with a single highly accumulated siRNA peak, in which only a 7-bp mutation in the predicted key region resulted in the loss of gene silencing (Fig. 4h-l). A small-RNA-seq analysis confirmed that the mutation resulted in the loss of the accumulation of the targeted single siRNA (Fig. 4n-m).

**Figure 4.**
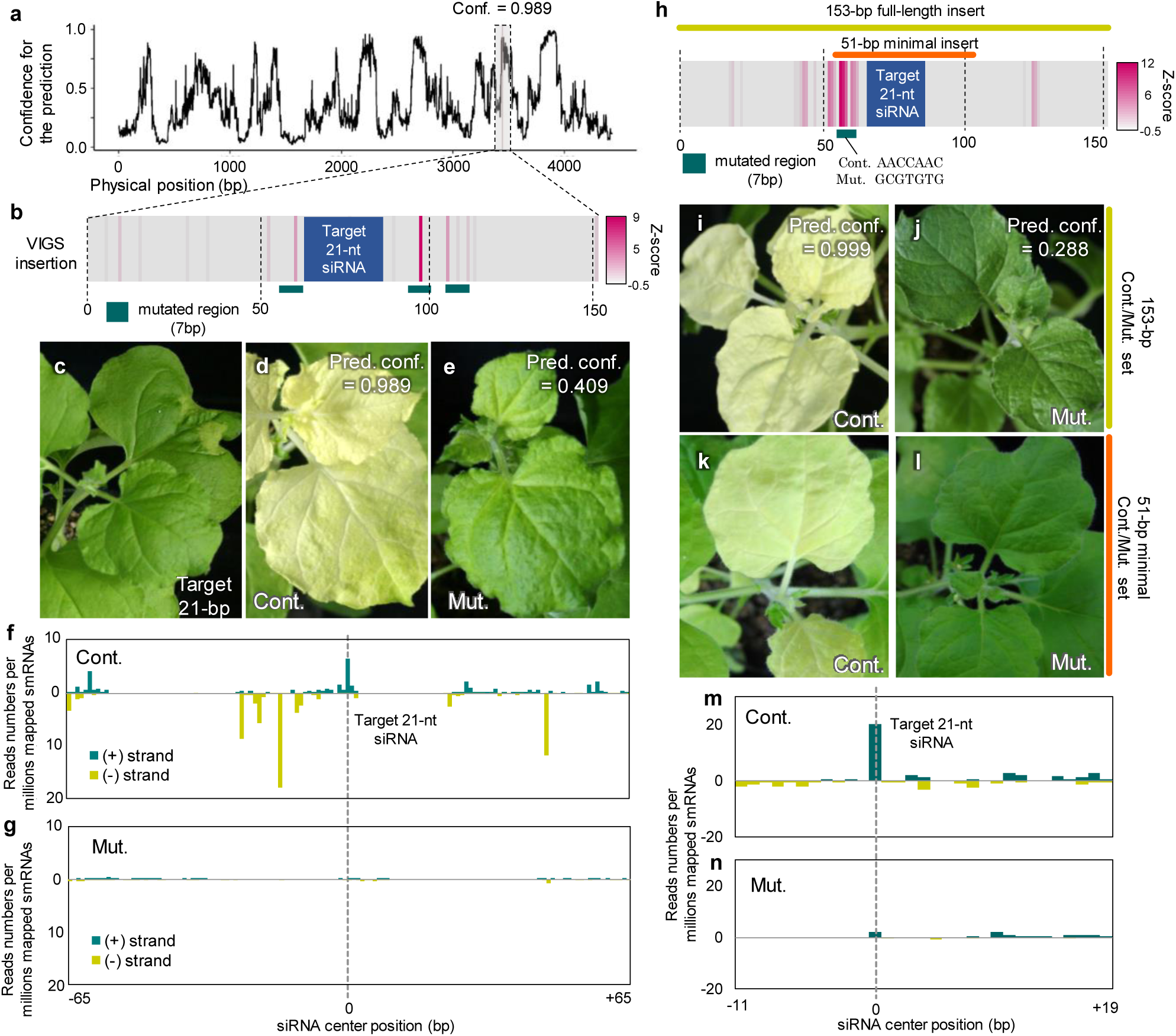
Experimental validation for the deep learning-guided estimation of the signal sequences required for siRNA accumulation. (**a**), Transition of the prediction confidence for siRNA accumulation of the *N. benthamiana* ChlH gene in the *C. sinensis* CvT model. (**b**), A 153-bp *ChlH* fragment predicted with high confidence to allow the accumulation of siRNA (conf. = 0.989). The standardized (Z-scored) attentions are shown in a heatmap. The residues with weighted attentions (or key sequences for the prediction) were artificially mutated, as indicated by green bars, to design the mutated fragment (Mut.). (**c-e**), Detection of leaf decoloration was used as an index for measuring effective RNAi, and the experiments were performed using VIGS and *ChlH*-inserted ALSV vectors. Introduction of only the accumulated 21-nt siRNA could not induce effective RNAi (**c**). The control (Cont.) 153-bp fragment insertion including the flanking regions with the key sequences (see panel **b**) induced clear decoloration (**d**), whereas the described artificially mutated insertion (Mut.) lost RNAi ability (**e**). (**f- g**), siRNA accumulation in the Cont. (**f**) and Mut. (**g**) *ChlH* genes. In the Mut. sample, not only the target siRNA accumulation but also the surrounding highly accumulated siRNA peaks were lost. (**h**), A 153-bp *ChlH* fragment in which only a single mutation (green bar) resulted in loss of effective RNAi ability (**i-j**). The orange bar indicated the putative minimal insert to induce effective RNAi (**k**), in which a single mutation similarly caused the loss of RNAi ability (**l**). siRNA was highly accumulated only in the target siRNA site in the Cont. (**m**); but lost in the Mut. sample (**n**).

**Figure 5.**
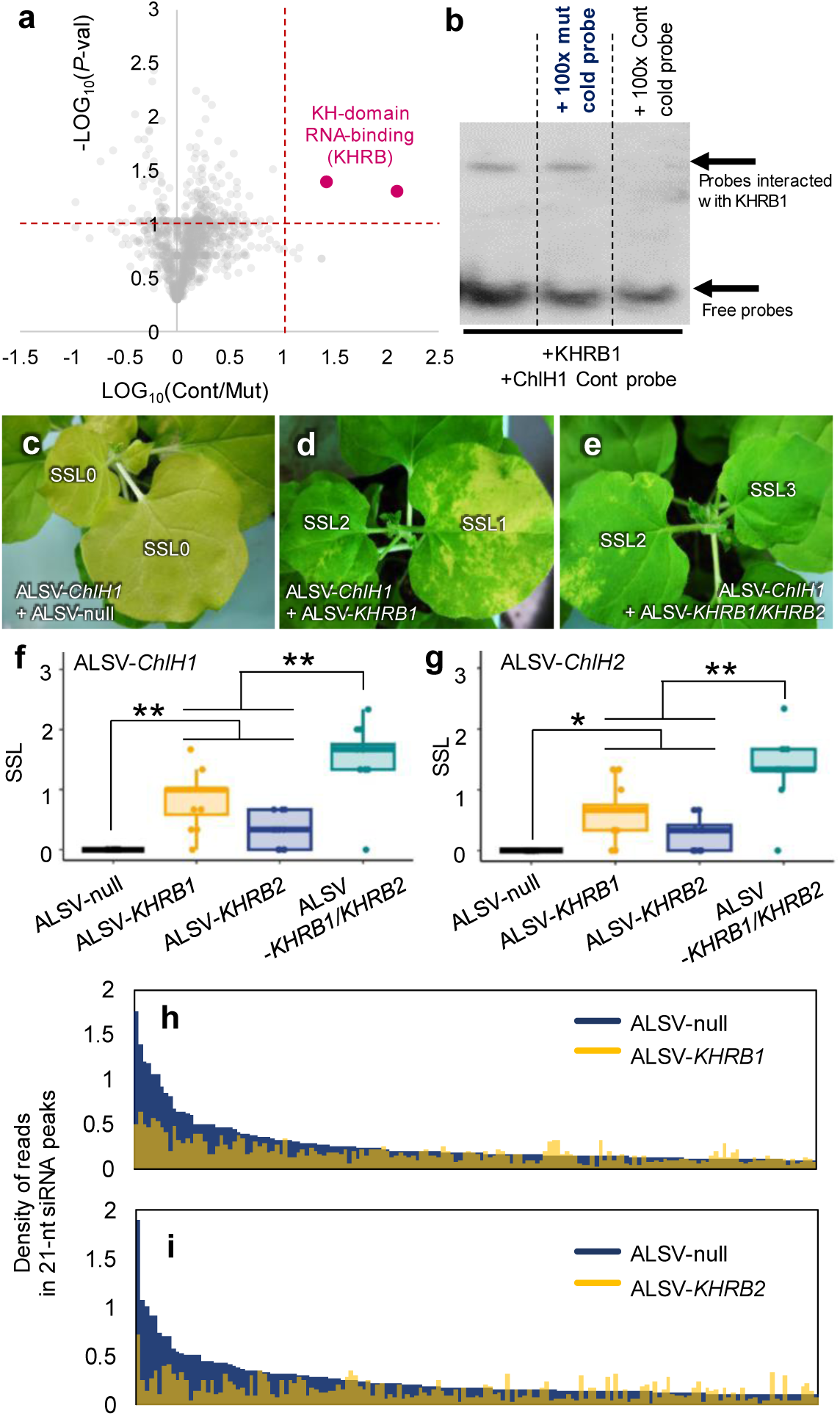
Novel KH-domain RNA-binding factors interact with the signal sequences for siRNA accumulation, which is required for effective RNAi. (**a**), Volcano plot of the 1,088 proteins that could potentially interact with the *ChlH* minimal insertions. Only two KH-domain RNA-binding (KHRB) proteins were detected using the criteria of *p* < 0.1 and >10 fold-change in Cont/Mut. (**b**), EMSA to confirm the direct interaction of the KHRB1 and the signal sequences. Addition of excess Cont. cold probe meant the binding signal was lost, whereas addition of excess Mut. cold probe did not affect the binding signal. (**c-e**), Suppression of *KHRB* genes expression resulted in gene silencing suppression in *N. benthamiana*. In contrast to the control (infection of non-inserted ALSV-null) (**c**), infection of the *KHRB1-* (**d**) or *KHRB/KHRB2*- inserted ALSV (**e**) suppresses the silencing effects (or leaf decoloration) by *ChlH* RNAi. SSL, silencing suppression level. SSL0-3 indicated 0-10%, 10-50%, 50-80%, and 80-100% greenish regions in a leaf, respectively. (**f-g**), Box plots of SSLs in ALSV with no insert (ALSV-null), ALSV- *KHRB1*, ALSV-*KHRB2*, and ALSV-*KHRB1/KHRB2* infected *N. benthamiana* leaves. The SSLs were detected using two independent *ChlH* fragments, *ChlH1* (Fig. 3h) and *ChlH2* (Fig. 3b). \**p* < 0.1, \*\**p* < 0.01 (one-sided Student’s *t*-test). (**h-i**), Genome-wide siRNA accumulation levels in highly accumulated siRNA peaks (21-nt read density >0.1%), in comparison to the control and *KHRB1-* or *KHRB2-*suppressed leaves in the identical *N. benthamiana* plants. Two *KHRB*s could independently regulate siRNA accumulation, especially when there are higher siRNA peaks.

The minimal *ChlH* insertion (defined as *ChlH1*) control/mutation (Cont./Mut.) set described above was used to identify factors recognizing the key region for siRNA accumulation. We used an RNA-centric approach to identify proteins associated with *ChlH1* RNA. We immobilized biotin-labeled *ChlH1* cont or mut RNAs on beads and applied crude proteins extracted from *N. benthamiana*, followed by on-bead digestion and analysis using nano LC-MS/MS (see Methods for the details). Enrichment analysis of the proteins that interacted with *ChlH1*-cont or *ChlH1*-mut (Fig. 4a, *N* = 1,088) suggested that only two novel proteins nested into the same family, K-homology (KH)-domain RNA-binding factor^27^, and primarily interacted only with *ChlH1*-cont RNA (p-value < 0.1, and fold-change > 10). The KH-domain RNA-binding protein family was first identified as the human hnRNP K protein^28^, and it is the second most frequently found family in the RNA-binding domain proteins^27^. Electrophoretic mobility shift assay (EMSA) analysis with biotin-labeled *ChlH1*- cont RNA probe and excess cold (non-labeled) cont/mut competitive RNA confirmed that the identified KH-domain proteins directly recognized the key region of the *ChlH1* RNAs (Fig. 4b). Although some KH-domain proteins were identified by relating them to various phenotypes in *Arabidopsis*^29–32^, the factors identified in this study nested into an undefined clade and were monophyletically conserved in a wide variety of land plant species, with some lineage-specific duplication events in angiosperms (Fig. S9). Induction of ALSV-mediated gene silencing by these two KH-domain RNA-binding factors, named *KHRB1* and *KHRB2* (Nbe.v.1.1.chr02g22450 and Nbe.v.1.1.chr08g23190, respectively, in the *N. benthamiana* genome), resulted in the fundamental suppression of decoloration caused by *ChlH1* RNAi in *N. benthamiana* leaves (Fig. 4c-g). Silencing of only *KHRB1* or *KHRB2* resulted in significant decoloration suppression effects (or silencing suppression level; SSL), and silencing both of them resulted in an even stronger SSL (Fig. 4e-g). Here, we infected the leaves with two independent *ChlH* insertions, *ChlH1* and *ChlH2* (used in Fig. 3b), and the suppression of *KHRB*s resulted in consistent SSLs between these two *ChlH* fragments (Fig. 4f and g). Genome-wide small-RNA-seq analyses indicated that the *KHRB1* or *KHRB2*- silenced *N. benthamiana* leaves exhibited substantially less siRNA accumulation, especially those in the category of highly accumulated siRNAs, than the control ALSV-null infected leaves (Fig. 4h-i). Together, these results indicated that the novel KH-domain RNA-binding factors identified in this study have the redundant function of recognizing the various key signal sequences that facilitate selective siRNA accumulation, which effectively results in RNAi.

In this study, our novel concept is an RNA-centric screening of RNA-protein interactions using key RNA sequences identified via Transformer-based explainable deep learning. So far, key RNA sequences for selective siRNA accumulation have not been defined, presumably because of their linguistic and positional flexibility. Here, Transformer-based deep learning allowed the definition (or visualization) of this flexible context to use further screening experiments, resulting in the identification of novel interactive frames between two KH-domain proteins and key RNA signal sequences for selective RNA accumulation. Although the KH-domain family is thought to be involved in pre-mRNA processing, RNA splicing, maintenance, and decay^24,26,28,30^, its ability to directly recognize RNAs has not been well-reported, especially in plants. In *Arabidopsis*, a few studies reported interactions between a KH-domain and other proteins, which depended on their phenotypic context^26,28^. Hence, this novel insight would be widely applicable to the regulation of various gene expression patterns in an siRNA site-specific manner. After successfully predicting a signal sequence site for high siRNA accumulation via our explainable-AI-based methods, a combination with gene-editing to make point mutations on the signal sequence site or to make a new signal sequence site would allow fine-tuning of specific siRNA regulation. This information would not only provide new insights in basic RNAi mechanism in plant biology but would also have various applications in agronomy or new-style crop breeding.

## Materials and Methods

### Mining of small-RNA-seq datasets

We downloaded the small-RNA-seq datasets (from the SRA-NCBI database, https://www.ncbi.nlm.nih.gov/sra) in 15 plant species covering a wide variety of land plant species, of which the details were given in Table S1. The 21-nt sequences were extracted from the fastq data, and mapped to each plant reference genome (downloaded from Phytozome13, https://phytozome-next.jgi.doe.gov/, see details in Table S1) with the Burrows-Wheeler aligner (BWA) mem (version 0.7.15) (http://bio-bwa.sourceforge.net/)^34^ allowing no mismatches (-n 0, in bwa-aln command), to characterize the 21-nt siRNA abundancy per 1-kb bin. Single siRNAs with a coverage > 20 were categorized as “highly accumulated” (or positive), while singlet (coverage = 1) siRNAs were categorized as negative samples. The positive/negative 21-nt siRNA nucleotides and their flanking genomic sequences (0, 15, 40, 60, and 90 bp) in each mapped genome were sense(+)/antisense(-) strand-specifically converted into the one-hot arrays with 4-channels, in which, namely A, T (U), G, and C are [1,0,0,0], [0,1,0,0], [0,0,1,0] and [0,0,0,1].

### Deep learning models

The following four model structures were used in the prediction of positive/negative samples: (1) a neural network consisting of four fully connected layers (Fully Connected Neural Network; FC), (2) a VGG-like 1-dimensional convolutional neural network (VGG-like 1D-CNN) modified from the previous report^19^, (3) a single-layer Transformer encoder mainly consisting of the self-attention mechanism proposed in Vaswani et al. 2017^10^, and (4) a CvT consisting of the described 1D-CNN and the 6-layer Transformer encoders (Fig. S3). The convolution mechanism captures the relationships of surrounding sequences, while the self-attention mechanism captures the relationships of distant positions. By combining CNN and Transformer encoder, we can leverage their respective strengths and compensate for their weaknesses. Additionally, the attention map calculated in the Transformer encoder indicates the weights of the sequences contributing to the prediction, providing high explainability for the inference. All model parameters were optimized with the cross-entropy loss function. We used Adam^35^ as the optimizer. The learning rate and number of epochs were turned 0.1-0.0001 and 5-100, respectively. Seventy-five percent of all sequences were used as training data, and the remaining 25% as test/validation data. The Receiver Operating Characteristic (ROC) curve was plotted based on the true positive-false positive rate of the predictions to calculate ROC area under curve (ROC-AUC) values as the index of prediction performance. Four-fold cross-validation was performed for all models to compare the prediction performance amongst the learning models. All procedures were run on Ubuntu 18.04 (Newtech, Cloudy-DP, 48 GB RAM for GPU). All of the applied codes described above have been released at https://github.com/matsuo-shinnosuke/siRNA_accum.

To investigate attributes in the actual and predicted positive siRNA, we classified the actual and predicted positive sequences of *O. sativa* (*N* = 5,663 for the actual positive and 694 for the predicted positive with prediction confidence > 0.9) into canonical genes, each transposable element (TE) family, and micro RNA (miRNA) using BLASTN. The reference sequences for TEs were derived from the rice genome project (https://rapdb.dna.affrc.go.jp/download/irgsp1.html), and for miRNAs, from rice miRNA information on mirBase (https://mirbase.org/). Fisher’s exact test was conducted to test the biases of attributes between the actual and predicted positive siRNA sequences in each TE and miRNA category.

### Visualization of the weighted residues for deep learning predictions

For the 1D-CNN models, we performed guided backpropagation^36^ on 151 bp (21bp + 65bp x 2) sequences predicted with high prediction confidence for the positive (conf. ≥ 0.9), according to the previous analysis^19^. Similarly, for sequences of 151 bp selected under the same conditions from the model created with CvT, we visualized the residues relevant to the positive prediction (conf. > 0.9) by outputting the attention in the encoder^10^. The codes used for visualizing these features are deposited at https://github.com/matsuo-shinnosuke/siRNA_accum. After standardizing the relevance level within the sequences, we performed hierarchical clustering based on relevance rates per position, which was visualized with heatmaps.

### Motif enrichment analyses

In the positive sequence fragments with a prediction confidence >0.9, each 10-bp flanking region of the residues with the highest attention (or total 21-bp) was collected in 5 plant species (*O. sativa*, *C. sinensis*, *V. vinifera*, *M. domestica*, and *A. arguta*), to detect potentially enriched nucleotide patterns in the high attention regions. With position-weight matrix (PWM)-based methods in the MEME suite (https://meme-suite.org/meme/), gap-free and gap-included motifs were explored with MEME^7^ (https://meme-suite.org/meme/tools/meme) and GLAM2scan^24^ (https://meme-suite.org/meme/tools/glam2scan), respectively. The explainability of the total significant motif (position *p-value* < 0.0001), with e-value < 10, was counted by using MAST^37^ (https://meme-suite.org/meme/tools/mast).

### VIGS induction with apple latent spherical viruses (ALSV)

We extracted total RNA from *Nicotiana benthamiana* young leaves using the PureLink® Plant RNA Reagent (refambion), and amplified 153-bp *ChlH* (AF014052.1), *PDS* (EU165355.1), *AGO1* (DQ321488.1), or *PHAN* (FR878011.1) gene fragments from the cDNAs, which were predicted as positive (or negative) with high prediction confidence (>0.95) in the CvT or 1d-CNN models trained with siRNA in some species. For *ChlH* gene, we ordered some synthesized DNA fragments (Eurofins Japan), in which the residues with weighted attention were artificially mutated (see Table SXX for the primer and synthesized fragment sequences). The 153-bp fragments were inserted into the multi-cloning site of the pCAM-ALSV2^38^, which encodes Apple Latent Spherical Virus (ALSV)-RNA2 (Li et al., 2004), according to the previous study^38^. Briefly, the control or mutated inserts amplified with adapter-included primers (Table S5) using PrimeSTAR GXL (TaKaRa, Japan), were purified by using the illustra™ GFX PCR DNA and Gel Band Purification Kit, followed by cloning into the pCAM-ALSV2 using the In-Fusion® HD Cloning Kit. The resultant pCAM-ALSV2 including an insert, pCAM-ALSV1, and a silencing suppressor p19-drived construct were transformed into Agrobacterium (strain EHA105) by electroporation, according to the previous study^38^. The cultured three Agrobacterium suspensions, of which the concentration was adjusted to OD600 = 1.0, were mixed in a 1:1:1 ratio, and infiltrated into young 3 leaves of *N. benthamiana* at the stage with 8-12 leaves. Approximately 3-4 weeks after the inoculation of transformed ALSV, 3 top leaves were used for detection of decoloration levels (or other phenotypic changes). Simultaneously, they were sampled and frozen in liquid nitrogen, followed by extraction of total RNA. From them, cDNA was synthesized using ReverTra Ace® qPCR RT Master Mix (Toyobo, Japan), and conduct PCR analysis to check the infection status of the transformed ALSVs.

### Small-RNA-seq analysis

Small-RNA seq analyses were conducted basically according to the previous study by Akagi et al. 2014^39^. Total RNAs were extracted from *N. benthamiana* leaves with PureLink™ Plant RNA Reagent (Thermo Fisher Scientific). Small RNA fraction was enriched by using the *mir*Vana™ miRNA Isolation Kit (Thermo Fisher Scientific), followed by preparation of small RNA libraries using the NEBNext Small RNA Library Prep Set for Illumina (NEB), according to the manufacture’s instruction. All of Illumina sequencing (PE150) was performed with NovaSeq6000 or NovaSeqX in Macrogen Japan. The paired sequences files were sorted with the index sequences, followed by trimmings of adapter sequences, low-quality reads (Phred score < 20), N-containing sequences, and reads ≥ 36 bases. The resultant small-RNA libraries were mapped to the *N. benthamiana* reference whole genes v1.1 (https://nbenthamiana.jp/)^40^, with the Burrows-Wheeler aligner (BWA), bwa-aln, (version 0.7.15) (http://bio-bwa.sourceforge.net/), allowing no mismatches (or -n = 0). From the generated sam-format files, information of 21-bp length reads was extracted to further analyze their locations and coverages.

### Affinity purification of proteins associated with *ChlH1* RNA

The 3’ biotinylated *ChlH* minimal Cont./Mut. RNA fragments (Fig. 3h) were synthesized by Hokkaido System Science Co., Ltd. Total 5 nmol of biotinylated RNAs mixed with 50 μL of Dynabeads M-280 streptavidin (Invitrogen) were washed with Citrate-Phosphate buffer (pH 5.0) for 3 times, followed by cross-linking with 20 mM dimethyl pimelimidate in 0.2 M triethanolamine (pH 8.2). A young *N. benthamiana* leaf was crushed under liquid nitrogen to extract crude proteins with 500 μL of 50 mM Tris-HCl (pH 7.5), including 150 mM NaCl, complete protease inhibitors mix cocktail tablet (Roche), and 1% Triton X-100, at 4°C for 30 min. The 200 μL supernatant was incubated with the cross-linked RNA-beads at 4°C for 120 min to allow RNA-protein complexes to form. After capturing the target proteins, beads were washed with wash buffer (50 mM Tris-HCl, 150 mM NaCl, 0.1% Triton X-100), and two more times with wash buffer excluding the detergent (50 mM Tris-HCl, 150 mM NaCl).

### Protein digestion and nano LC-MS/MS data acquisition

Captured proteins are digested on the beads. The streptavidin beads were suspended in 20 μl of digestion buffer (8 M urea, 250 mM Tris-HCl [pH 8.5]), reduced with 25 mM tris(2- carboxyethyl)phosphine (TCEP) at 37°C for 15 min and alkylated using 25 mM iodoacetamide at 37°C for 30 min in the dark, both with shaking at 1,200 rpm following the standard protocol. Proteins on beads were directly digested with 0.1 μg Lys-C (FUJIFILM Wako, Japan) at 37°C for 3 h with shaking at 1,000 rpm. After dilution to a urea concentration of 2 M with 50 mM Tris-HCl (pH 8.5) followed by addition of 1 mM CaCl_2_, Lys-C digest was further digested with 0.1 μg trypsin (Promega) with shaking at 1,000 rpm at 37°C overnight. The digestion was stopped by adding 5 μl of 20% TFA, and then Lys-C/trypsin–double digested peptides were desalted using a GL-Tip SDB (GL Science) according to the manufacturer’s instructions. Nano-LC-MS/MS analysis was performed using an EASY-nLC 1200 LC system (ThermoFisher Scientific) connected to an Orbitrap Exploris 480 hybrid quadrupole-orbitrap mass spectrometer (Thermo Fisher Scientific). Desalted samples were dissolved in 200 μl of 2% acetonitrile (0.1% TFA), and 7.5-μl aliquots of the supernatant were used for LC-MS/MS analysis. Samples were loaded in direct injection mode and separated on a nano-HPLC capillary column (Aurora column [75 μm I.D. × 250 mm], IonOpticks) with a gradient of 3– 32% acetonitrile (containing 0.1% formic acid) over 90 min at a flow rate of 300 nl/min. The Orbitrap Exploris 480 mass spectrometer was operated in data-dependent acquisition mode with dynamic exclusion enabled (20 s). MS/MS scans were performed by higher-energy collisional dissociation with the normalized collision energy set to 30. The MS/MS raw files were processed and analyzed with Proteome Discoverer 2.5 (Thermo Fisher Scientific) using the SEQUEST HT algorithm, searching against the Nbe.v1 *N. benthamiana* protein database.

### Electrophoretic mobility shift assay

To confirm the direct interaction of the *ChlH* minimal RNA fragments and KHRB1 protein, we conducted electrophoretic mobility shift assay (EMSA). The full length of KHRB1 CDS was amplified with adapter-included primers (Table S5) and PrimeSTAR GXL (TaKaRa) to be subcloned into pENTR vector (Invitrogen), and then cloned into pDEST15 with LR clonase, to designate pDEST15-KHRB1. The pDEST15-KHRB1 was transformed to BL21 *E. coli* which produces GST- KHRB1 fusion protein. Modifying to the previous study^38^, GST-KHRB1 fusion protein was extracted with using GST Fusion Protein Purification Kit (Funakoshi). The GST-KHRB1 protein and Cont. biotinylated *ChlH* RNA as the signal probe were incubated with non-labeled Cont. and Mut. ChlH RNAs as the cold probes at room temperature for 30min. The mixtures were run in 10% polyacrylamide gel, to transfer to Biodyne plus (Pall). The transferred biotinylated RNAs were hybridized with Streptavidin Alkaline Phosphatase (Promega), to detect by chemiluminescence method with CDP-star (Sigma-Aldrich).

### Evolutionary analysis

The orthologs of the two KH-domain RNA-binding (KHRB) proteins identified in *N. benthamiana* (Nbe.v.1.1.chr02g22450 and Nbe.v.1.1.chr08g23190) were detected in *Arabidopsis thaliana*, *Populus trichocarpa*, *Vitis vinifera*, *Solanum lycopersicum*, *Oryza sativa*, and outgroup land plants species, *Selaginella moellendorffii* and *Marchantia polymorpha*, using BLASTp (<1e-20) in Phytozome (JGI release version 13.0, https://phytozome.jgi.doe.gov/pz/portal.html). Protein sequences were aligned using MAFFT ver. 7 with the L-INS-i model, followed by automatic trimming procedures with TrimAl v1.2^42^, followed by calculation of the evolutionary topology with iqtree (-m MFP -bb 1000 -nt AUTO)^43^. Bootstraps were shown on the branches as 1/10 of the calculated values.

## Data Availability Statement

All Illumina sequencing (transcriptome and DNA methylome) data have been deposited in the DDBJ database: Short Read Archives (SRA) database (BioProject ID PRJDB18427, Biosample ID SAMD00799105-SAMD00799112). The raw mass spectrometry data have been deposited in the ProteomeXchange Consortium via the jPOST partner repository under accession number PXD053741. The source data underlying Fig. 4a and Supplementary Table. S4 is provided as a Source Data file.

## ACKNOWLEDGEMENT

We thank Dr. Reina Komiya (Okinawa Institute of Science and Technology) for the discussion and comments and Edanz (https://jp.edanz.com/ac) for editing a draft of this manuscript. This work was supported by PRESTO from Japan Science and Technology Agency (JST) [JPMJPR20D1], Grant-in-Aid for Transformative Research Areas (A) from JSPS [22H05172 and 22H05173] to T.A. and S.U. and JSPS Research Fellow from JSPS [23KJ1612] to E.K.

## AUTHOR CONTRIBUTION

S.U. and T.A. conceived the study. Y.M., S.U., and T.A. designed experiments. N.E., E.K., S.M., N.F., S.N., and T.A. conducted the experiments. N.E., S.M., S.N., and T.A. analyzed the data. N.E., and T.A. drafted the manuscript. All authors approved the manuscript.

